# TranceptEVE: Combining Family-specific and Family-agnostic Models of Protein Sequences for Improved Fitness Prediction

**DOI:** 10.1101/2022.12.07.519495

**Authors:** Pascal Notin, Lood Van Niekerk, Aaron W Kollasch, Daniel Ritter, Yarin Gal, Debora S. Marks

## Abstract

Modeling the fitness landscape of protein sequences has historically relied on training models on family-specific sets of homologous sequences called Multiple Sequence Alignments. Many proteins are however difficult to align or have shallow alignments which limits the potential scope of alignment-based methods. Not subject to these limitations, large protein language models trained on non-aligned sequences across protein families have achieved increasingly high predictive performance – but have not yet fully bridged the gap with their alignment-based counterparts. In this work, we introduce TranceptEVE – a hybrid method between family-specific and family-agnostic models that seeks to build on the relative strengths from each approach. Our method gracefully adapts to the depth of the alignment, fully relying on its autoregressive transformer when dealing with shallow alignments and leaning more heavily on the family-specific models for proteins with deeper alignments. Besides its broader application scope, it achieves state-of-the-art performance for mutation effects prediction, both in terms of correlation with experimental assays and with clinical annotations from ClinVar.

## 1 Introduction

Modeling the fitness landscape of proteins has historically relied on multiple-sequence alignments (MSAs) – sets of homologous sequences that are all aligned in the same position coordinate system. Models trained on MSAs have progressively increased in expressiveness – from models relying on position-specific features in the early days [Ng and Henikoff, 2001], to models relying on pairs of residues Hopf et al. [2017], and ultimately on full sequences [Riesselman et al., 2018, Frazer et al., 2021, Shin et al., 2021]. However, family-specific alignment-based models require sufficiently deep alignments to learn the rich distributions that would capture the complex dependencies across protein residues, and there are many proteins (e.g., disordered proteins) that are difficult to align or for which alignments are relatively shallow. Models trained across protein families (e.g., Alley et al. [2019]) have been introduced as a potential solution for that limitation, relying on the hypothesis that biochemical constraints that are learned for certain protein families or domains (e.g., with many homologs) can be transferred to similar proteins or domains with shallower alignments. Building on recent progress in Natural Language Processing, large-scale transformer models trained large quantities of unaligned sequences across families have made tremendous progress in that direction Rives et al. [2021], Elnaggar et al. [2021], Meier et al. [2021], Madani et al. [2020], Nijkamp et al. [2022], Ferruz et al. [2022], Hesslow et al. [2022] – yet, none of these models has been able so far to bridge the gap with alignment-based architectures in the zero-shot setting. To achieve comparable performance, alignments still have to be used for predictions, giving rise to ‘hybrid models’ that borrow strengths from family-specific and family-agnostic approaches: Unirep [Alley et al., 2019] and ESM-1v [Meier et al., 2021] rely on fine-tuning pretrained models on an MSA for the protein of interest; the MSA Transformer [Rao et al., 2021] learns a model of multiple-sequence alignments across thousands of different families; and, more recently, Tranception [Notin et al., 2022] achieves state-of-the-art fitness prediction performance by combining predictions from a large autoregressive transformer with the retrieval of an MSA at inference time. In this work, we introduce TranceptEVE, a hybrid model building on the concepts developed in Tranception. In addition to extracting residue-specific distributions from the retrieved MSA, we learn an EVE model Frazer et al. [2021] over that MSA to capture potentially critical epistatic effects. This leads to superior fitness prediction performance when comparing with experimental results from Deep Mutational Scanning (DMS) assays (§ 3.1). Furthermore, since the aggregation coefficients between the autoregressive transformer and the EVE model are dependent on the depth of the retrieved MSA, our approach gracefully adapts to all types of proteins: if the protein has few homologs, we mainly rely on the autoregressive transformer; if the retrieved MSA is deep, we lean more heavily on the family-specific EVE model. Lastly, we demonstrate that this approach outperforms all prior methods when predicting the effects of mutations on human proteins based on clinical labels (§ 3.2). Our contributions are as follows:

- We develop TranceptEVE, a model combining a family-agnostic Tranception model with family-specific EVE models for superior fitness prediction performance (§ 2);
- We evaluate TranceptEVE against the 94 Deep Mutational Scanning (DMS) assays from the ProteinGym benchmarks [Notin et al., 2022], extending the set of baselines (e.g., Progen2, RITA, Unirep, ESM-1b) previously reported (§3.1);
- We demonstrate the ability of TranceptEVE to correctly predict the effects of genetic mutations in humans based on expert-labelled variants from ClinVar (§ 3.2).

## 2 TranceptEVE

We are interested in assessing the fitness of mutated proteins, i.e., their ability to perform their function. The distribution of protein sequences observed in nature is the result of billions of evolutionary experiments that selected out the unfit variants. By learning a distribution over these sequences we implicitly capture the biochemical constraints that characterize fit variants. In practice, we train a generative model to learn that distribution and then assess the relative fitness of a mutated protein compared to that of a wild-type sequence from the corresponding family via their log likelihood ratio under the learned model:

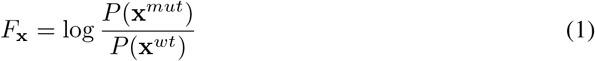

For a protein x composed of residues (*x*_1_, *x*_2_,&, *x_L_*), the likelihood factorizes via the chain rule as:

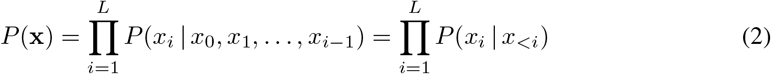

In TranceptEVE, the log-likelihood at each position log *P*(*x_i_*| *x_<i_*) is obtained as a weighted arithmetic average between the log-likelihoods from two distinct models: 1) log *P_T_*(*x_i_*| *x_<i_*) from Tranception [Notin et al., 2022], a large autoregressive transformer model trained across protein families and 2) log *P_E_*(*x_i_*| *x_<i_*) from EVE [Frazer et al., 2021], a variational autoencoder trained on a retrieved MSA for the protein family of interest. More precisely, after training the protein-specific EVE model, we obtain log *P_E_* (*x_i_*| *x_<i_*) by inputting into the VAE the wild-type sequence from which the MSA was acquired and then averaging the resulting log-softmax outputs from the decoder network across a large number of samples from the approximate posterior *z* ~ *q*(*z* | x^*wt*^) and from the distribution over decoder parameters. As such, the decoder output from the EVE model acts as a constant family-specific prior distribution over amino acids at each sequence position that is independent from the particular protein sequence we wish to estimate the fitness of: log *P_E_* (*x_i_* | *x_<i_*) = log *P_E_* (*x_i_*). Besides preserving autoregressiveness – which is desirable for novel proteins generation, this approach presents several practical advantages over naive ensembling (§ B.6) and, critically, it allows to handle the scoring of insertions and deletions (§ B.3).

We thus estimate each residue probability in Eq. 2 via the following expression:

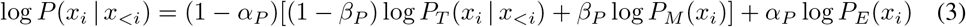

where log *P_M_* is the empirical distribution over amino acids at each position of the retrieved MSA (i.e., the inner bracket corresponds to Tranception with MSA retrieval); *α_P_* and *β_P_* are constants that solely depends on the depth of the corresponding alignment: when the alignment is shallow we rely fully on the autoregressive predictions log *P_T_*, and when the MSA is deeper we lean more heavily on the MSA and EVE log priors (log *P_M_* and log *P_E_* resp., § B.2). In practice, we score sequences from both directions (N→C and C→N) for increased fitness prediction performance (see details in § B.1).

## 3 Experiments

### 3.1 Correlation with Deep Mutational Scanning experiments

#### ProteinGym

ProteinGym Notin et al. [2022] is the largest collection of Deep Mutational Scanning (DMS) assays for assessing mutation effects predictors. It consists of two different benchmarks measuring mutations made via *substitutions* (~1.5M missense variants across 87 DMS assays) and *indels* (~300K mutants across 7 DMS assays). In our experiments, we compared against all baselines already available in ProteinGym (e.g., Tranception, EVE, MSA Transformer) and added the following: Unirep [Alley et al., 2019], RITA [Hesslow et al., 2022], Progen2 [Nijkamp et al., 2022] and ESM-1b (original model from Rives et al. [2021] with extensions from Brandes et al. [2022]; Appendix C.1).

#### Results

TranceptEVE outperforms all other baselines in the ProteinGym benchmark in aggregate performance (Spearman or AUC) and for each MSA depth class (Table 1; on par with ESM-1v on the high MSA depth subset). It also performs best for each taxon (human and other eukaryotes, prokaryotes and viruses – see Table 4), for each mutation depth (Table 5) and on the indels benchmark (Table 6).

**Table 1:**
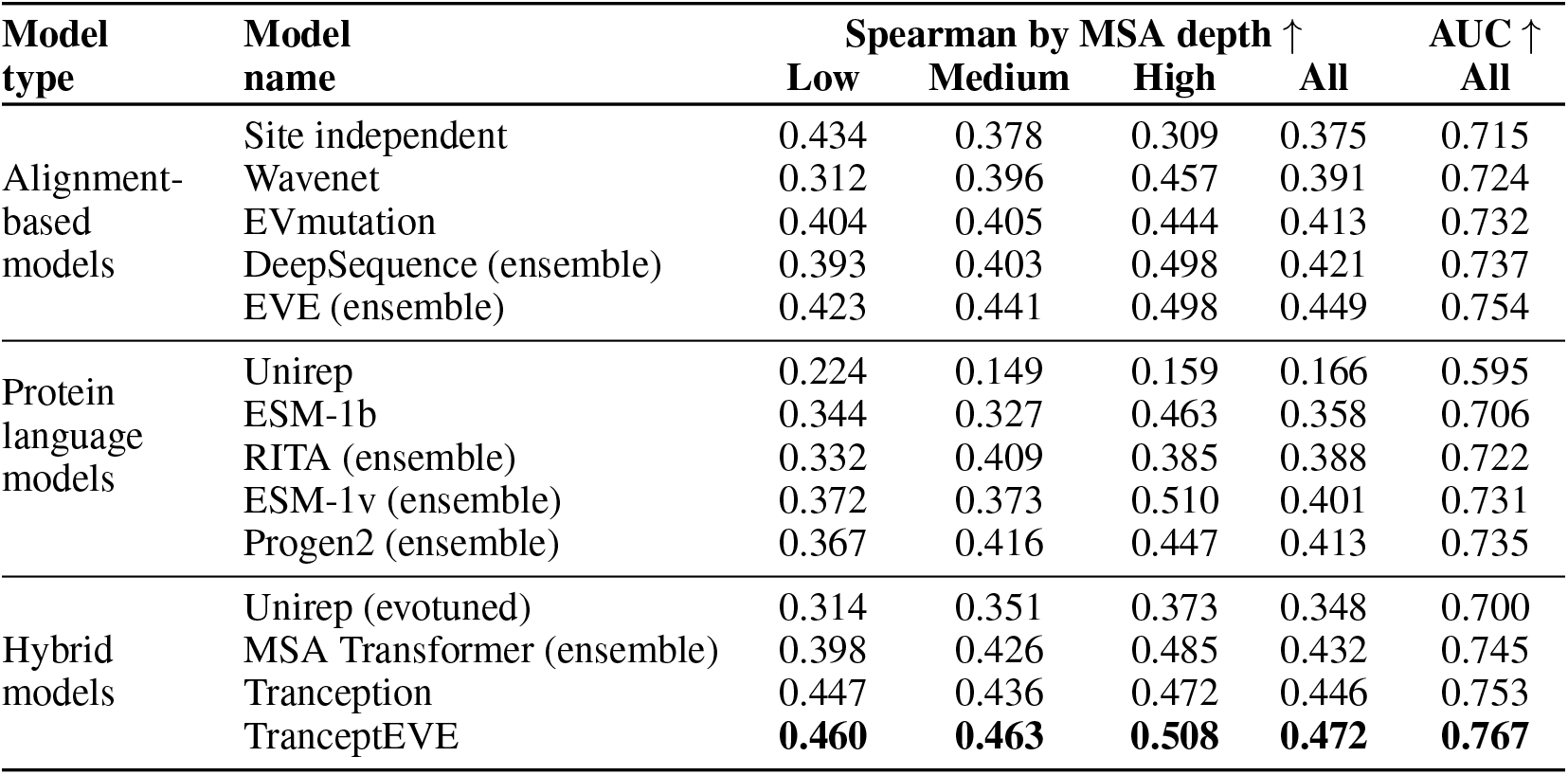
Average AUC and Spearman’s rank correlation between model scores and experimental measurements by MSA depth on the ProteinGym substitution benchmark. Alignment depth is measured by the ratio of the effective number of sequences *N*_eff_ in the MSA, following Hopf et al. [2017], by the length covered *L* (Low: *N*_eff_/*L* <1; Medium: 1 < *N*_eff_/*L* <100; High: *N*_eff_/*L* >100). Statistical significance analysis reported in Table 3 in Appendix.

### 3.2 Predicting the effects of human genetic variation on disease risk

#### ClinVar ClinVar

[Landrum et al., 2015] is a database of genetic variants for human genes and the associated interpretations of clinical significance for reported conditions. We focused on the same ~3k disease genes as in Frazer et al. [2021] and evaluated the ability of mutation effects predictors to predict the pathogenicity of genetic variants (~31k pathogenic and benign ClinVar classifications).

#### Results

The average protein-level AUC of TranceptEVE across all proteins is markedly higher than that of EVE [Frazer et al., 2021], ESM-1b [Brandes et al., 2022] and all other supervised or unsupervised baselines (Fig. 2). TranceptEVE also has a higher spearman correlation with DMS assays of human proteins – whether we consider the subset from Frazer et al. [2021] as in Fig. 2 or all DMS assays for human proteins in ProteinGym (Appendix C.2).

**Figure 1:**
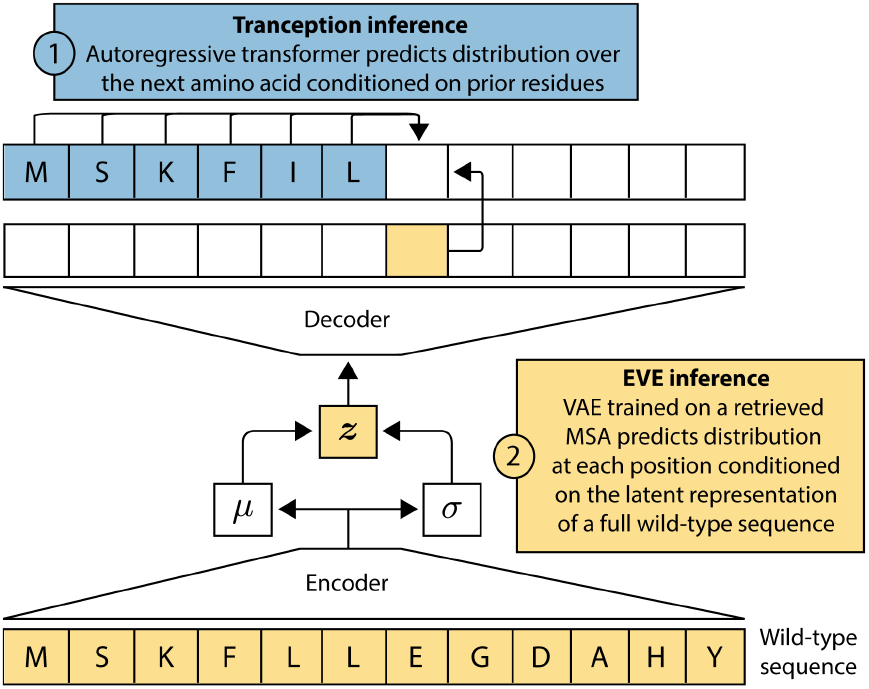
TranceptEVE combines a family-agnostic autoregressive transformer and a family-specific EVE model at inference. TranceptEVE predictions are based on two complementary modes of inference: 1) an autoregressive model – Tranception – makes predictions at a given residue based on the context of previous amino acids in the sequence; 2) a proteinspecific variational autoencoder – EVE – learns a distribution over amino acids at each position based on dependencies across the full sequence observed in a retrieved Multiple Sequence Alignment.

**Figure 2:**
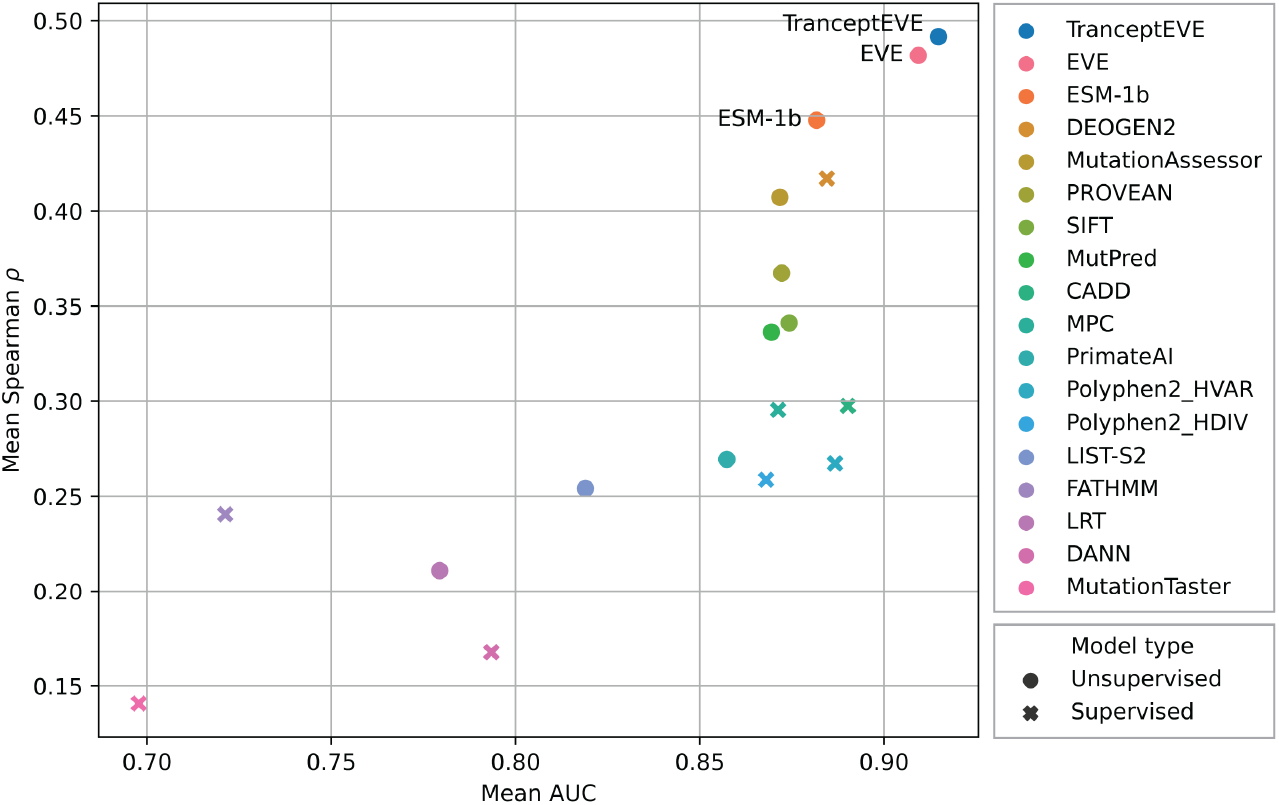
Performance comparison of TranceptEVE to state-of-the-art computational variant effect predictors: nine unsupervised and eight supervised. Performance estimated against known clinical labels (avg. AUC over disease genes in ClinVar (x axis)), and high-throughput functional assays developed to assess the clinical effect of variants (avg. Spearman’s rank correlation (y axis)).

## 4 Conclusion

We have introduced TranceptEVE, a hybrid between the family-agnostic Tranception model and the family-specific EVE model. TranceptEVE outperforms all baselines both in terms of correlation with DMS experiments from ProteinGym and with clinical annotations from ClinVar. This performance improvement holds across alignment depths, taxa and mutation depths. Future work will investigate how to learn family-agnostic and family-specific models in a unified architecture trained end-to-end.

## Appendix

## A Background

Large-scale models with architectures derived from Natural Language Processing have displayed increasing performance for protein fitness prediction, particularly in the zero-shot setting [Alley et al., 2019, Meier et al., 2021, Hesslow et al., 2022]. Despite this, these methods are most effective when they incorporate information specific to the target protein family. Most recently, Tranception combined predictions from a large autoregressive transformer trained across protein families with predictions based on retrieval from an MSA at inference time. Retrieval predictions are based on the empirical distribution over amino acid at each position, akin to the way they are derived in a site-independent model [Ng and Henikoff, 2001]. Higher-order models such as DeepSequence or EVE [Riesselman et al., 2018, Frazer et al., 2021] consistently outperform site-independent models when trained on the same alignments. However, the combination of higher-order alignment models with alignment-free, family-agnostic models such as Tranception for unsupervised fitness prediction has not yet been described.

## B Modeling details

### B.1 Inference details

We chose Tranception and EVE as they are respectively the best family-agnostic and family-specific mutation effect predictors at the time of this writing (based on correlation with experimental results from the ProteinGym benchmarks). Furthermore, in early experiments, these two models showed the best complementary when performing naive ensembling between models (testing all possible pairs from baselines in [Notin et al., 2022]).

We obtain the ‘EVE log prior’ as follows:

- We train an EVE model (or, in the case of the ProteinGym analyses, an ensemble of 5 EVE models with different random initializations) on an MSA retrieved from a wild type sequence that is representative of the protein family of interest (e.g., canonical sequence in Uniprot);
- We encode that same wild type sequence in the latent space of the trained bayesian VAE from EVE, and then decode it a large number of times (200k samples);
- We average the resulting log softmax probabilities (i.e., the decoder outputs) across samples;
- If we have trained an ensemble of 5 EVE models, we also average across the models in the ensemble;
- The resulting tensor represents a log probability over the amino acid vocabulary as each position in the sequence (i.e., log *P_E_*(*x*)), akin to the empirical distributions obtained in the MSA retrieval of Tranception (i.e., log *P_M_* (*x*)).

As discussed in § 2, we score sequences from both directions (i.e., N→C and C→N), then take the arithmetic average of each log probability:

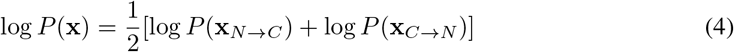

### B.2 Aggregation coefficients based on MSA depth

Instead of the constant aggregation coefficient used in Tranception, we use aggregation coefficients (i.e., α and β in Equation 2) which depends on the depth of the retrieved MSA for a given protein family (see Table 2). This enables TranceptEVE to seamlessly adapt to all possible protein families by leaning more heavily on the most relevant mode of inference depending on the situation: if the protein family of interest has no or very few homologs, we rely exclusively on the autoregressive transformer; if the MSA is deeper, we give more importance to the MSA and EVE log priors.

**Table 2:** Aggregation coefficient for EVE log prior (α) and MSA log prior (β) based on MSA depth.

|  | MSA depth (Nb. sequences) |  |  |  |  |
| --- | --- | --- | --- | --- | --- |
|  | < 10 | < 10<sup>2</sup> | < 10<sup>3</sup> | < 10<sup>5</sup> | ≥ 10<sup>5</sup> |
| $\alpha$ | 0.0 | 0.3 | 0.6 | 0.7 | 0.8 |
| $\beta$ | 0.0 | 0.1 | 0.3 | 0.4 | 0.5 |

### B.3 Indels scoring details

When scoring insertions & deletions (‘indels’), we adapt the log prior distributions by re-aligning the protein to be scored to the family-specific MSA: deletions in the protein sequence translate to deletions of the corresponding positions in the priors; insertions lead to the addition of dummy columns at these positions in the priors, which are then ignored in the final weighted average (i.e., in Equation 2) such that the predictions at these positions rely solely on the autoregressive transformer.

### B.4 Recalibrating models probabilities

Since the transformer in Tranception and the VAE in EVE are trained independently on different data domains, their output probabilities are not identically calibrated. After factoring in the relative weights with respect to MSA depth as discussed in Appendix B.2, we would like each model to have the same overall importance in the predictions from Equation 2. We thus iteratively recalibrate the EVE log probabilities via temperature scaling so that the two models have the same mean output log softmax when given the wild type sequence as input.

### B.5 Scope of the EVE log prior

In the original EVE architecture [Frazer et al., 2021], we were only leveraging positions that were sufficiently-covered in the MSA. As a result, the corresponding EVE models were only able to score mutations at these well-covered positions. There is however no strict constraint to model only well-covered positions, and we have observed in practice that also including the non well-covered positions in the EVE models was 1) not detrimental to the predictive performance when scoring mutations at well-covered positions only (e.g., same overall performance on the ProteinGym benchmark) 2) would allow to score mutations at non well-covered positions as well. In this work, the reported performance for EVE models on the ProteinGym benchmark (e.g., Table 1) is obtained when modeling all positions and scoring all possible mutants from the benchmark. TranceptEVE fully adapts to the characteristics of the underlying EVE model: if the EVE model was trained on well-covered positions only, it bases its predictions for the non well-covered positions on the autoregressive transformer and retrieved MSA only (as in Tranception with retrieval); otherwise it leverages the autoregressive transformer and the two log priors.

### B.6 Comparison with standard model ensembling

The aggregation described in Equation 2 can be perceived as performing some form of model ensembling between Tranception and EVE (we do not perform a strict ‘model ensembling’ since the EVE log prior is independent from the particular sequences to score and is instead protein-family specific). While we observe comparable overall performance between our TranceptEVE scheme and a standard ensembling of Tranception and EVE (e.g., average of standardized log likelihood ratios from Tranception and standardized delta ELBOs from EVE) on the ProteinGym substitution benchmark (eg., same average Spearman and AUC), our proposed scheme however presents several significant advantages:

- The TranceptEVE scoring scheme preserves autoregressiveness during decoding, which allows it to support novel sequence generation for protein design (unlike with standard ensembling where we only obtain a single score for the full sequence with the delta ELBOs output by EVE);
- TranceptEVE can be used to score indels as per the extensions described in Appendix B.3 (unlike naive ensembling since EVE is unable to score insertions or deletions given its fixed-size MLP encoder);
- Standard ensembling of Tranception and EVE requires the scores to be standard scaled before averaging (since the Tranception and EVE models scores are on different scales). This operation requires scores from both models for a representative set of sequences (e.g., scores from all singles) to compute the relevant score statistics (i.e., mean and variance). It is not clear which representative set to use in practice (e.g., should we use the same set for mutations at different mutation depths?) and represents a potentially large computational overhead. TranceptEVE is not impacted by any of these issues;
- In TraanceptEVE we only score the wild type sequence with EVE and use the same EVE log prior to score all mutated sequences of interest. With standard ensembling we need to score each sequence with both Tranception and EVE.

## C Additional experimental results

### C.1 Correlation with Deep Mutational Scanning experiments

#### Baselines

For all baselines already present in Notin et al. [2022], we use the same model implementations and scores. These baselines are as follows: Site independent model and EVmutation [Hopf et al., 2017], DeepSequence [Riesselman et al., 2018], Wavenet [Shin et al., 2021], EVE [Frazer et al., 2021], ESM-1v [Meier et al., 2021], MSA Transformer [Rao et al., 2021] and Tranception [Notin et al., 2022]. For the new baselines, we use the official github repositories for the implementations of Unirep [Alley et al., 2019], RITA [Hesslow et al., 2022] and Progen2 [Nijkamp et al., 2022]. Evo-tuning for Unirep is performed on the relevant protein MSAs from ProteinGym. For all RITA and Progen2 model variants, we score sequences from both directions (similar to what is described in Appendix B.1 for TranceptEVE). For these two baselines, the ensemble version is obtained by ensembling the different sizes of the corresponding model (e.g., RITA S, RITA M, RITA L and RITA XL). For ESM-1b, we use the official ESM repository for model checkpoint and mutation effects predictions. We further extend the codebase to handle sequences longer than the model context size (i.e., longer than 1,022 amino acids) as per the procedure described in Brandes et al. [2022].

#### Statistical significance

Given the high correlation between baselines across assays (e.g., certain assays are difficult to predict for all models, others easy for all models), we assess the statistical significance of the results reported in Table 1 via the bootstrap standard deviation of the *difference* between TranceptEVE and other baselines (Table 3).

#### Additional results

We further compare the fitness prediction performance of TranceptEVE and all baselines on the ProteinGym substitution benchmark by taxon (Table 4) and by mutation depth (Table 5). Table 6 provides a performance summary on the ProteinGym indels benchmark.

### C.2 Predicting the effects of human genetic variation on disease risk

#### Baselines

Besides TranceptEVE and ESM-1b, all baselines reported in Fig 2 are taken directly from Frazer et al. [2021]: DEOGEN2 [Raimondi et al., 2017], MutationAssessor [Reva et al., 2011], PROVEAN [Choi et al., 2012], SIFT [Sim et al., 2012], MutPred [Li et al., 2009], CADD [Rentzsch et al., 2019], MPC [Samocha et al., 2017], PrimateAI [Sundaram et al., 2018], Polyphen2 [Adzhubei et al., 2010], LIST-S2 [Malhis et al., 2020], FATHMM [Shihab et al., 2013], LRT [Chun and Fay, 2009], DANN [Quang et al., 2015] and MutationTaster [Schwarz et al., 2010]. For ESM-1b, the performance on ClinVar (avg. AUC) is obtained directly from the mutation predictions provided in Brandes et al. [2022]. The scoring of the EVE DMS assays (y-axis in Fig 2) is obtained with the same codebase as described in Appendix C.1 for the ProteinGym experiments. For TranceptEVE, we use the model checkpoints from Frazer et al. [2021] (single seed model only).

#### Results

As in Frazer et al. [2021], we assess model performance on the ClinVar benchmark in terms of the average per-protein AUC since: 1) it is consistent with the way these models are mainly used in practice (i.e., identify which mutants *for a given gene* are likely pathogenic for *a specific disease*) 2) it is consistent with the way we look at performance on DMS assays (ie., average of per-assay performance) and 3) reporting an ‘overall AUC’ (i.e., grouping together all available labels from ClinVar) would confuse together unrelated pathologies (e.g., with unrelated disease mechanisms, of different severity, with different difficulty of predictability). We further deep dive into the top 3 models as per Fig. 2 in Table C.2. TranceptEVE outperforms ESM-1b and EVE on all benchmarks.

**Table 3:** Average Spearman’s rank correlation between model scores and experimental measurements on the ProteinGym substitution benchmark, and standard error of the difference to TranceptEVE. The standard error reported is the non-parametric bootstrap standard error of the difference between the Spearman performance of TranceptEVE and that of a given baseline, computed over 10k bootstrap samples from the set of proteins in the ProteinGym substitution benchmark.

| Model type | Model name | Average Spearman | Standard error of diff. to TranceptEVE |
| --- | --- | --- | --- |
| Alignment-based models | Site independent | 0.375 | 0.012 |
|  | Wavenet | 0.391 | 0.010 |
|  | EVmutation | 0.413 | 0.005 |
|  | DeepSequence (ensemble) | 0.421 | 0.009 |
|  | EVE (ensemble) | 0.449 | 0.005 |
| Protein language models | Unirep | 0.166 | 0.024 |
|  | ESM-1b | 0.358 | 0.019 |
|  | RITA (ensemble) | 0.388 | 0.014 |
|  | ESM-1v (ensemble) | 0.401 | 0.019 |
|  | Progen2 (ensemble) | 0.413 | 0.011 |
| Hybrid models | Unirep (evotuned) | 0.348 | 0.014 |
|  | MSA Transformer (ensemble) | 0.432 | 0.012 |
|  | Tranception (w/ retrieval) | 0.446 | 0.005 |
|  | TranceptEVE | **0.472** | - |

**Table 4:** Average Spearman’s rank correlation *ρ* between model scores and experimental measurements by taxon.

| Model type | Model name | Spearman by Taxon ↑ | | | | |
| --- | --- | --- | --- | --- | --- | --- |
|  |  | Human | Other eukar. | Prokaryote | Virus | All |
| Alignment-based models | Site independent | 0.366 | 0.417 | 0.324 | 0.412 | 0.375 |
|  | Wavenet | 0.378 | 0.446 | 0.472 | 0.308 | 0.391 |
|  | EVmutation | 0.385 | 0.448 | 0.472 | 0.381 | 0.413 |
|  | DeepSequence (ens.) | 0.400 | 0.493 | 0.492 | 0.348 | 0.421 |
|  | EVE (ens.) | 0.408 | 0.499 | 0.500 | 0.435 | 0.449 |
| Protein language models | Unirep | 0.254 | 0.226 | 0.166 | 0.01 | 0.166 |
|  | ESM-1b | 0.391 | 0.442 | 0.485 | 0.153 | 0.358 |
|  | RITA (ens.) | 0.378 | 0.387 | 0.381 | 0.410 | 0.388 |
|  | ESM-1v (ens.) | 0.425 | 0.441 | 0.502 | 0.257 | 0.401 |
|  | Progen2 (ens.) | 0.399 | 0.459 | 0.462 | 0.364 | 0.413 |
| Hybrid models | Unirep (evotuned) | 0.334 | 0.338 | 0.372 | 0.351 | 0.348 |
|  | MSA Transformer (ens.) | 0.394 | 0.487 | 0.498 | 0.398 | 0.432 |
|  | Tranception | 0.417 | 0.503 | 0.478 | 0.429 | 0.446 |
|  | TranceptEVE | **0.440** | **0.518** | **0.514** | **0.454** | **0.472** |

**Table 5:** Average Spearman’s rank correlation *ρ* between model scores and experimental measurements by mutation depth.

| Model type | Model name | Spearman by mutation depth ↑ | | | | | |
| --- | --- | --- | --- | --- | --- | --- | --- |
|  |  | 1 | 2 | 3 | 4 | 5+ | All |
| Alignment-based models | Site independent | 0.375 | 0.322 | 0.285 | 0.321 | 0.407 | 0.375 |
|  | Wavenet | 0.389 | 0.344 | 0.324 | 0.282 | 0.35 | 0.391 |
|  | EVmutation | 0.414 | 0.401 | 0.403 | 0.335 | 0.427 | 0.413 |
|  | DeepSequence (ens.) | 0.419 | 0.394 | 0.407 | 0.353 | 0.436 | 0.421 |
|  | EVE (ens.) | 0.45 | 0.409 | 0.405 | 0.351 | 0.429 | 0.449 |
| Protein language models | Unirep | 0.161 | 0.201 | 0.097 | 0.126 | 0.192 | 0.166 |
|  | ESM-1b | 0.351 | 0.318 | 0.215 | 0.161 | 0.318 | 0.358 |
|  | RITA (ens.) | 0.380 | 0.236 | 0.135 | 0.181 | 0.246 | 0.388 |
|  | ESM-1v (ens.) | 0.400 | 0.309 | 0.203 | 0.165 | 0.253 | 0.401 |
|  | Progen2 (ens.) | 0.410 | 0.312 | 0.233 | 0.228 | 0.282 | 0.413 |
| Hybrid models | Unirep (evotuned) | 0.339 | 0.284 | 0.279 | 0.246 | 0.323 | 0.348 |
|  | MSA Transformer (ens.) | 0.433 | 0.374 | 0.401 | 0.351 | 0.418 | 0.432 |
|  | Tranception | 0.442 | 0.427 | 0.440 | 0.370 | 0.461 | 0.446 |
|  | TranceptEVE | **0.472** | **0.444** | **0.456** | **0.389** | **0.464** | **0.472** |

**Table 6:** Average AUC and Spearman’s rank correlation between model scores and experimental measurements on the ProteinGym indel benchmark.

| Model name | Spearman ↑ | AUC ↑ |
| --- | --- | --- |
| Progen2 (ensemble) | 0.407 | 0.748 |
| Wavenet | 0.412 | 0.740 |
| RITA (ensemble) | 0.446 | 0.761 |
| Tranception (Large) | 0.464 | 0.773 |
| Tranception (Medium) | 0.509 | 0.796 |
| TranceptEVE (Large) | 0.466 | 0.774 |
| TranceptEVE (Medium) | **0.516** | **0.799** |

**Table 7:** Performance of top mutation effect predictors on different human proteins fitness benchmarks. The ClinVar and EVE DMS set benchmarks are based on Frazer et al. [2021]. The last benchmark is obtained by filtering the assays in the ProteinGym substitution benchmark [Notin et al., 2022] to only keep human protein assays.

| Dataset | Metric | ESM-1b | EVE | TranceptEVE |
| --- | --- | --- | --- | --- |
| ClinVar (3k disease-related genes) | Avg. AUC | 0.882 | 0.909 | **0.915** |
| EVE DMS set (11 assays) | Avg. Spearman | 0.448 | 0.482 | **0.492** |
| ProteinGym human proteins set (34 assays) | Avg. Spearman | 0.391 | 0.408 | **0.440** |

## References

I. A. Adzhubei, S. Schmidt, L. Peshkin, V. E. Ramensky, A. Gerasimova, P. Bork, A. S. Kondrashov, and S. R. Sunyaev. A method and server for predicting damaging missense mutations. Nature Methods, 7(4):248–249, Apr. 2010. ISSN 1548-7091, 1548-7105. doi:10.1038/nmeth0410-248. URL http://www.nature.com/articles/nmeth0410-248.

E. C. Alley, G. Khimulya, S. Biswas, M. AlQuraishi, and G. M. Church. Unified rational protein engineering with sequence-based deep representation learning. Nature Methods, pages 1–8, 2019.

N. Brandes, G. Goldman, C. H. Wang, C. J. Ye, and V. Ntranos. Genome-wide prediction of disease variants with a deep protein language model. bioRxiv, 2022.

Y. Choi, G. E. Sims, S. Murphy, J. R. Miller, and A. P. Chan. Predicting the functional effect of amino acid substitutions and indels. 2012.

S. Chun and J. C. Fay. Identification of deleterious mutations within three human genomes. Genome research, 19(9):1553–1561, 2009.

A. Elnaggar, M. Heinzinger, C. Dallago, G. Rehawi, W. Yu, L. Jones, T. Gibbs, T. B. Fehér, C. Angerer, M. Steinegger, D. Bhowmik, and B. Rost. Prottrans: Towards cracking the language of lifes code through self-supervised deep learning and high performance computing. IEEE transactions on pattern analysis and machine intelligence, PP, 2021.

N. Ferruz, S. Schmidt, and B. Höcker. Protgpt2 is a deep unsupervised language model for protein design. Nature Communications, 13, 2022.

J. Frazer, P. Notin, M. Dias, A. Gomez, J. K. Min, K. P. Brock, Y. Gal, and D. S. Marks. Disease variant prediction with deep generative models of evolutionary data. Nature, 2021.

D. Hesslow, N. ed. Zanichelli, P. Notin, I. Poli, and D. S. Marks. Rita: a study on scaling up generative protein sequence models. ArXiv, abs/2205.05789, 2022.

T. A. Hopf, J. B. Ingraham, F. J. Poelwijk, C. P. Schärfe, M. Springer, C. Sander, and D. S. Marks. Mutation effects predicted from sequence co-variation. Nature biotechnology, 35(2):128–135, 2017.

M. J. Landrum, J. M. Lee, M. Benson, G. Brown, C. Chao, S. Chitipiralla, B. Gu, J. Hart, D. Hoffman, J. Hoover, W. Jang, K. Katz, M. Ovetsky, G. Riley, A. Sethi, R. Tully, R. Villamarin-Salomon, W. Rubinstein, and D. R. Maglott. ClinVar: public archive of interpretations of clinically relevant variants. Nucleic Acids Research, 44(D1):D862–D868, 11 2015. ISSN 0305-1048. doi:10.1093/nar/gkv1222. URL https://doi.org/10.1093/nar/gkv1222.

B. Li, V. G. Krishnan, M. E. Mort, F. Xin, K. K. Kamati, D. N. Cooper, S. D. Mooney, and P. Radi-vojac. Automated inference of molecular mechanisms of disease from amino acid substitutions. Bioinformatics, 25(21):2744–2750, 2009.

A. Madani, B. McCann, N. Naik, N. S. Keskar, N. Anand, R. R. Eguchi, P.-S. Huang, and R. Socher. Progen: Language modeling for protein generation, 2020.

N. Malhis, M. Jacobson, S. J. Jones, and J. Gsponer. List-s2: taxonomy based sorting of deleterious missense mutations across species. Nucleic acids research, 48(W1):W154–W161, 2020.

J. Meier, R. Rao, R. Verkuil, J. Liu, T. Sercu, and A. Rives. Language models enable zero-shot prediction of the effects of mutations on protein function. bioRxiv, 2021. doi:10.1101/2021.07.09.450648. URL https://www.biorxiv.org/content/early/2021/07/10/2021.07.09.450648.

P. C. Ng and S. Henikoff. Predicting deleterious amino acid substitutions. Genome Research, 11(5): 863–874, 2001. doi:10.1101/gr.176601. URL http://genome.cshlp.org/content/11/5/863.abstract.

E. Nijkamp, J. A. Ruffolo, E. N. Weinstein, N. Naik, and A. Madani. Progen2: Exploring the boundaries of protein language models. ArXiv, abs/2206.13517, 2022.

P. Notin, M. Dias, J. Frazer, J. Marchena-Hurtado, A. N. Gomez, D. S. Marks, and Y. Gal. Tranception: protein fitness prediction with autoregressive transformers and inference-time retrieval. In ICML, 2022.

D. Quang, Y. Chen, and X. Xie. Dann: a deep learning approach for annotating the pathogenicity of genetic variants. Bioinformatics, 31(5):761–763, 2015.

D. Raimondi, I. Tanyalcin, J. Ferté, A. Gazzo, G. Orlando, T. Lenaerts, M. Rooman, and W. Vranken. Deogen2: prediction and interactive visualization of single amino acid variant deleteriousness in human proteins. Nucleic acids research, 45(W1):W201–W206, 2017.

R. Rao, J. Liu, R. Verkuil, J. Meier, J. F. Canny, P. Abbeel, T. Sercu, and A. Rives. Msa transformer. bioRxiv, 2021. doi:10.1101/2021.02.12.430858. URL https://www.biorxiv.org/content/early/2021/02/13/2021.02.12.430858.

P. Rentzsch, D. Witten, G. M. Cooper, J. Shendure, and M. Kircher. Cadd: predicting the deleteriousness of variants throughout the human genome. Nucleic acids research, 47(D1):D886–D894, 2019.

B. Reva, Y. Antipin, and C. Sander. Predicting the functional impact of protein mutations: application to cancer genomics. Nucleic acids research, 39(17):e118–e118, 2011.

A. J. Riesselman, J. B. Ingraham, and D. S. Marks. Deep generative models of genetic variation capture the effects of mutations. Nature methods, 15(10):816–822, 2018.

A. Rives, J. Meier, T. Sercu, S. Goyal, Z. Lin, J. Liu, D. Guo, M. Ott, C. L. Zitnick, J. Ma, et al. Biological structure and function emerge from scaling unsupervised learning to 250 million protein sequences. Proceedings of the National Academy of Sciences, 118(15), 2021.

K. E. Samocha, J. A. Kosmicki, K. J. Karczewski, A. H. O’Donnell-Luria, E. Pierce-Hoffman, D. G. MacArthur, B. M. Neale, and M. J. Daly. Regional missense constraint improves variant deleteriousness prediction. BioRxiv, page 148353, 2017.

J. M. Schwarz, C. Rödelsperger, M. Schuelke, and D. Seelow. Mutationtaster evaluates diseasecausing potential of sequence alterations. Nature methods, 7(8):575–576, 2010.

H. A. Shihab, J. Gough, D. N. Cooper, P. D. Stenson, G. L. Barker, K. J. Edwards, I. N. Day, and T. R. Gaunt. Predicting the functional, molecular, and phenotypic consequences of amino acid substitutions using hidden markov models. Human mutation, 34(1):57–65, 2013.

J.-E. Shin, A. J. Riesselman, A. W. Kollasch, C. McMahon, E. Simon, C. Sander, A. Manglik, A. C. Kruse, and D. S. Marks. Protein design and variant prediction using autoregressive generative models. Nature communications, 12(1):1–11, 2021.

N.-L. Sim, P. Kumar, J. Hu, S. Henikoff, G. Schneider, and P. C. Ng. SIFT web server: predicting effects of amino acid substitutions on proteins. Nucleic Acids Research, 40(W1):W452–W457, July 2012. ISSN 0305-1048, 1362-4962. doi:10.1093/nar/gks539. URL https://academic.oup.com/nar/article-lookup/doi/10.1093/nar/gks539.

L. Sundaram, H. Gao, S. R. Padigepati, J. F. McRae, Y. Li, J. A. Kosmicki, N. Fritzilas, J. Hakenberg, A. Dutta, J. Shon, et al. Predicting the clinical impact of human mutation with deep neural networks. Nature genetics, 50(8):1161–1170, 2018.

